# FlexiBAC: a versatile, open-source baculovirus vector system for protein expression, secretion, and proteolytic processing

**DOI:** 10.1101/470955

**Authors:** Régis P. Lemaitre, Aliona Bogdanova, Barbara Borgonovo, Jeffrey B. Woodruff, David N. Drechsel

## Abstract

Baculovirus-mediated expression in insect cells is a powerful approach for protein production. However, many existing methods are time consuming, offer limited options for protein tagging, and are unsuitable for secreted proteins requiring proteolytic maturation, such as TGF-β family growth factors. To overcome these limitations, we engineered “FlexiBAC”, a system that simplifies baculovirus production and permits furin-driven proteolytic maturation of targets. This system allows recombinant baculovirus formation inside insect cells and reduces the time between initial cloning and protein production to 13 days. FlexiBAC includes 146 shuttle vectors that append combinations of purification tags, fluorescent markers, proteolytic cleavage sites, trafficking signals, and chemical conjugation tags to the termini of the target protein. We demonstrate that this system can be used to produce high levels of mature, active forms of TGF-β family growth factors, such as Activin A, as well as other proteins that are typically difficult to reconstitute, such as proteins rich in coiled-coil, low complexity, and disordered domains.

## INTRODUCTION

Protein production using baculovirus-infected insect cells combines biological safety with high-level expression of functional proteins. Eukaryotic proteins produced recombinantly in this system generally adopt a native, folded conformation and contain the appropriate post-translational modifications. The large baculovirus genome (ranging between 80 and 180 kbp, depending on the species) can accommodate large insertions, making it ideal for expressing large heteromeric protein complexes. Moreover, insect cells can grow in serum-free media, which greatly facilitates purification of secreted proteins from the conditioned media.

Despite these advantages, recombinant protein production using baculovirus is time consuming. The time between initial cloning and protein expression lasted 3 to 4 weeks in the earliest baculovirus expression vector systems that relied on plaque selection of recombinants into the polyhedron locus after homologous recombination in insect cells (Jarvis, 2009; van Oers et al., 2015). To make baculovirus production simpler and faster, more recent protocols have facilitated production of recombinant baculovirus genomes using site-specific transposition in *E. coli* (Bac-to-Bac®, Thermo Fisher Scientific) or homologous recombination in insect cells (flashBAC™, Oxford Expression Technologies). Both systems employ the polyhedrin promoter to drive high-level, late expression of the target protein(s) of interest. While the former system is more popular, the latter system is faster, as it reduces the window between initial cloning and protein expression to ~14 days. One major drawback of these commercial systems is that they only offer limited options for tagging the protein of interest: 6xHis, GST, and S-tag options are available, but fluorophores or tags for conjugating chemical probes are not. Purification of difficult proteins would be greatly facilitated by a baculovirus vector system that offers a larger combination of purification and solubility tags. Furthermore, as reconstituted systems are analyzed more and more via light microscopy, there is a pressing need for more fluorophore tagging options.

Most baculovirus expression systems are also not suited for high-level secretion of recombinant proteins. The secretory pathway in insect cells has limited capacity and cannot handle the efflux of recombinant proteins during baculovirus infection (Kaba et al., 2004). During late infection, baculovirus produce cathepsin and chitinase, two highly-expressed enzymes which are secreted by insect cells. In the wild, these enzymes are essential for liquefaction of the larval host, but they play no role in baculovirus replication in cultured cells. It was proposed that eliminating these proteins would reduce the viral protein load passing through the secretory pathway and would free up room for recombinant target proteins. Indeed, deletion of the genes encoding cathepsin and chitinase dramatically improved secretion of recombinant target proteins; unexpectedly, these mutations also reduced degradation of both secreted and non-secreted target proteins (Hitchman et al., 2010; Kaba et al., 2004). Thus, knocking out genes encoding cathepsin and chitinase should generally improve the efficiency and versatility of baculovirus expression systems.

Another limitation of baculoviral expression systems is that they are not equipped to handle high-volume post-translational processing of secreted proteins. Many secreted proteins require proteolytic trimming to reach a fully active, mature form. One prominent example is the transforming growth factor beta (TGF-β) member Activin A, which must be cleaved by furin convertases in the Golgi apparatus to become an active signaling ligand (Leduc et al., 1992). Baculovirus-mediated expression of Activin A largely resulted in the secretion of an inactive pro-form peptide (Cronin et al., 1998). Co-infection of baculoviruses expressing furin and Activin A greatly increased secretion of mature Activin A, indicating that furin activity is limited due to insufficient endogenous expression levels (Laprise et al., 1998). Engineering a single baculovirus to express furin in addition to the target gene of interest would increase yields of active TGF-β and other secreted peptides that require furin processing, while eliminating the need for co-infection.

To overcome these limitations, we designed an open source baculovirus expression system called FlexiBAC. Our system limits the time between initial cloning and protein isolation to 13 days and incorporates rescue of a doubly defective bacmid to ensure that all infective viruses contain the recombinant target gene. FlexiBAC includes a compatible, modular shuttle vector set (146 vectors) that appends combinations of various purification tags, fluorescent markers, proteolytic cleavage sites, trafficking signals, and chemical conjugation tags to the user’s target protein. To allow high yields of secreted proteins, the FlexiBAC viral genome includes deletions of genes encoding cathepsin and chitinase. We also present a specialized version of FlexiBAC that improves the maturation of secreted proteins requiring furin proteolysis. We demonstrate that FlexiBAC is suited for the expression of proteins deemed difficult to reconstitute, including mature TGF-β family growth factors, intrinsically disordered proteins, as well as proteins containing numerous coiled-coil domains.

## RESULTS

### Overview of the FlexiBAC system

To streamline the production of viral stocks, we engineered a bacmid encoding a replication-<u>def</u>ective <u>bac</u>ulovirus, which we term “DefBac” (Figure 1). This bacmid is designed to recombine with one of 146 compatible shuttle vectors, called pOCC vectors, where the target gene of interest (GOI) is inserted (Table 1 and 2). Transfection of *Spodoptera frugiperda* (Sf9) insect cells with linearized DefBac and a pOCC vector permits their recombination and produces infectious baculovirus. The GOI is then expressed late in the lytic cycle using the polydedrin promoter. Viruses are produced and escape the Sf9 cells only if the recombination between pOCC and DefBac was successful. One round of viral amplification is typically required before the virus is used to express the target protein (see Protocol section below). In the following sections, we describe construction of DefBac and the pOCC shuttle vectors.

**Figure 1.**
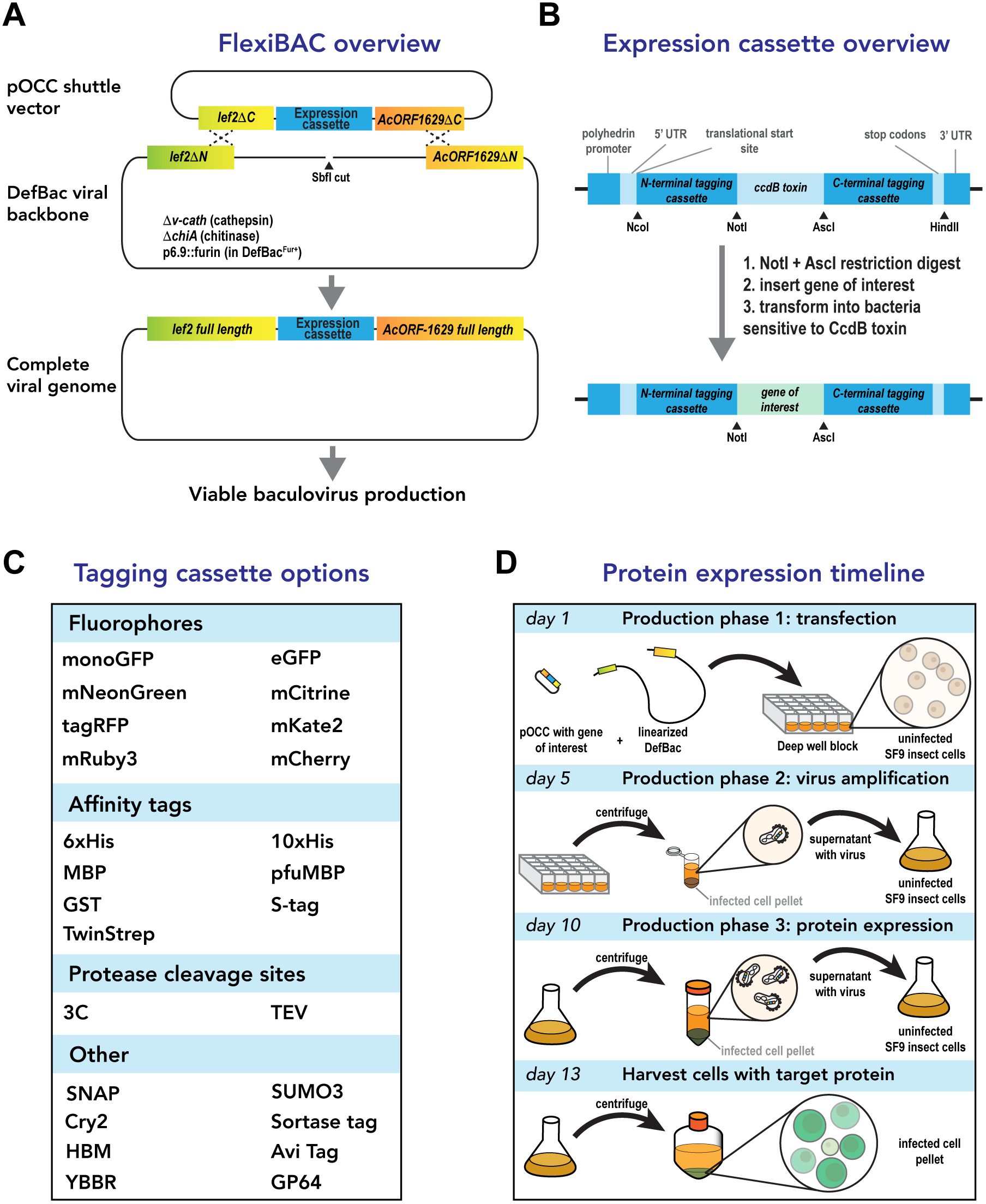
Overview of the FlexiBAC baculovirus expression system. A. Recombination between a SbfI-linearized defective viral backbone (“DefBac”) and a shuttle vector (“pOCC”, which contains the target gene of interest) creates a viral genome capable of producing infectious virus. Recombination occurs between complementary *lef2* and *AcORF1629* truncations located on the pOCC vector and the DefBac viral backbone. Recombination generates full-length versions of *lef2* and *AcORF1629* genes, which are needed for baculovirus production. No virus is produced without proper recombination, thus eliminating the need for post-production screening for recombinant virus. The DefBac viral backbone also included deletions in the genes encoding cathepsin and chitinase. A second version of DefBac, called DefBac^Fur+^, expresses the convertase furin. B. Each pOCC shuttle vector contains a modular expression cassette that allows insertion and swapping of the gene of interest, N-terminal tags, and C-terminal tags using classic cloning techniques (restriction sites are shown with arrowheads). A gene encoding the ccdB toxin selects against plasmids lacking the gene of interest. C. pOCC shuttle plasmids encode a variety of tags that can be appended to the target protein of interest. Tags are easily combined and swapped to create customizable N-terminal or C-terminal fusions. Descriptions of each tag and the available combinations (146) are shown in Table 1 and available upon request. The most commonly used plasmids (54) are shown in Table 2 and are available at www.addgene.org. D. Timeline for production of recombinant baculovirus using the FlexiBAC system. On day 1, the user transfects SF9 insect cells with pOCC shuttle vector (with target gene of interest) and linearized DefBac DNA. Within some insect cells, pOCC and DefBac will recombine and this event generates an infectious baculovirus that propagates throughout the culture. On day 5, the user collects the conditioned media (which contains released baculovirus) and uses it to infect fresh SF9 insect cells. On day 10, the user uses the conditioned media again to infect fresh SF9 cells. On day 13, the user harvests the infected SF9 cells containing the target protein of interest (represented by green, GFP+ cells). For secreted protein targets the conditioned medium is collected after this phase.

**Table 1.** Inventory of available pOCC shuttle vectors. The user’s gene of interest can be inserted into 146 available vectors using NotI and AscI restriction cloning. See associated .pdf file. Description of tags:

|  |  |
| --- | --- |
| 3C | human rhinovirus 3C protease cleavage site |
| GST | glutathione S-transferase |
| MBP | E.coli maltose-binding protein |
| PfuMBP | P. furiosus maltose-binding protein |
| HIS6 | hexahistidine |
| HIS10 | decahistidine |
| TwinStrep | WSHPQFEK-GGGSGGGSGG-SA-WSHPQFEK |
| Dendra2 | green-to-red photoswitchable fluorescent protein Dendra, derived from octocoral Dendronephthya sp. |
| LPETGG | sortase tag |
| mCherry | monomeric mutant fluorescent protein (FP) derived from drFP583 isolated from Discosoma |
| mKate2 | monomeric mutant FP derived from eqFP578 isolated from Entacmaea quadricolor |
| mNeonGreen | monomeric mutant FP derived from LanYFP isolated from Branchiostoma lanceolatum |
| monoGFP | monomeric mutant FP derived from Aequorea Victoria |
| mRuby3 | monomeric mutant FP derived from eqFP611 isolated from Entacmaea quadricolor |
| tagRFP | monomeric mutant FP derived from eqFP578 isolated from Entacmaea quadricolor |
| TEV | tobacco etch virus (TEV) protease |
| S-tag | peptide derived from pancreatic ribonuclease A |
| YBBR | peptide derived from B. subtilis ybbR |
| AviTag | 15 AA peptide tag (GLNDIFEAQKIEWHE) substrate for E. coli BirA enzyme |
| eGFP | enhance green fluorescent protein |
| GP64 | leader sequence for the glycoprotein 64 from GP64 of Autographa californicamultiple nucleopolyhedrovirus (AcMNPV) |
| HBM | leader sequence for honey bee mellitin |
| mCitrine | yellow mutant of GFP |
| GG | sortase tag |
| SUMO3 | Small ubiquitin-related modifier 3 |
| Cry2 | cryptochrome protein 2 from Arabidopsis thaliana (light switchable aggregation tag ) |

**Table 2.** Inventory of pOCC shuttle vectors available at www.addgene.org. The 54 most popular shuttle vectors were deposited to Addgene. See associated .pdf file.

#### A doubly defective viral backbone ensures production of complete recombinant viruses containing the gene of interest

One of the earliest, and still very popular, systems used to generate recombinant baculovirus for protein expression (Bac-to-Bac, Thermo Fisher) employs site-specific transposition of a baculovirus genome, propagated as a bacmid, with a cloning plasmid that contains the gene of interest. Transposition occurs in bacteria, which can take ~1 week to select and isolate positive clones. To streamline recombinant virus preparation, we modified the *Autographa californica* baculovirus bacmid (bMON14272) by deleting the 3’-ends of two essential genes: *lef-2* and *AcOrf-1629*. These genes flank the polyhedrin locus and are required for baculovirus production in cultured insect cells (Kitts and Possee, 1993) (Ho Je et al., 2001) (Zhao et al., 2003) (Wu et al., 2010). Cells transfected with this defective bacmid never appeared infected with baculovirus, and the cells continued to grow and divide at a rate similar to non-transfected controls (data not shown). In contrast, insect cells transfected with the unmodified parental bacmid produced infected cells 72 hours post transfection, as expected. We refer to this doubly defective bacmid as H092. The design ensures that the parental bacmid lacking the gene of interest will not produce functional baculovirus.

We designed H092 so that infectivity is restored by site-specific recombination with a pOCC shuttle vector, which contains the gene of interest flanked by 3’-segments of *lef-2* and *AcOrf-1629* that complement the truncations in H092 (Figure 1A). Recombination at the *lef-2* and *AcOrf-1629* loci restores functionality of these genes, thus producing infective, recombinant baculovirus genome that expresses the gene of interest (Figure 1A). Sf9 insect cells efficiently promote homologous recombination, suggesting that co-transfection of H092 and pOCC in Sf9 cells could reliably produce recombinant bacmid and functional virus. Indeed, co-transfection of H092 and a shuttle vector expressing GFP, under the control of the polyhedrin promoter, produced infected, GFP-positive Sf9 cells four days post-transfection (see Figure 2 for more examples).

**Figure 2.**
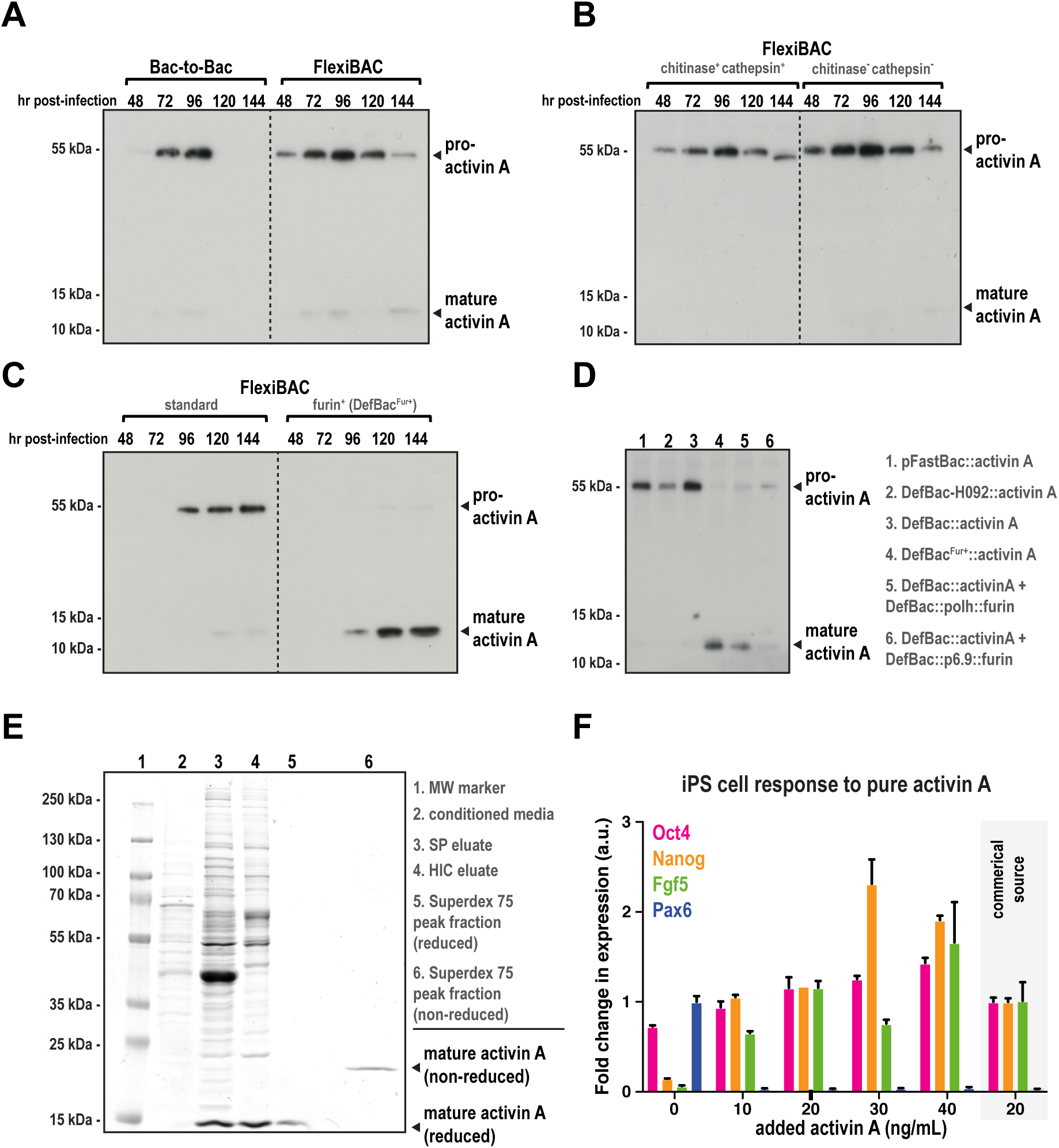
Efficient production of mature TGF-β member Activin A using FlexiBAC. A. SF9 insect cells were infected with recombinant baculovirus generated using the commercial Bac-to-Bac system (ThermoFisher) or the FlexiBAC system. Cell supernatants were collected at the indicated times after infection, resolved on 4-20% gradient SDS-PAGE gels under reducing conditions, then analyzed by western blot with anti-Activin A IgG. Bands corresponding to pro-Activin A (uncleaved) and mature Activin A (cleaved) are indicated. B. Activin A secretion from insect cells infected with baculovirus generated using the precursor DefBac backbone (H092) vs. the current DefBac backbone. The current system includes deletions in the genes encoding viral chitinase and cathepsin. C. Activin A maturation and secretion from insect cells infected with DefBac-derived baculovirus vs. DefBac^Fur+^-derived baculovirus. In addition to expressing the target protein of interest, DefBac^Fur+^ expresses furin convertase, which converts pro-Activin A to its mature form. D. Insect cells were either singly infected or co-infected with the following virus: pFastBac::Activin A (lane 1), DefBac-H092::Activin A (lane 2), DefBac::Activin A (lane 3), DefBac^Fur+^::Activin A (lane 4), DefBac::Activin A and DefBac::polh::Furin (lane 5), DefBac::Activin A and DefBac::p6.9::Furin (lane6). Each virus was added to insect cells at MOI = 0.2 except for DefBac::p6.9::Furin (MOI = 2). The conditioned media was analyzed by western blot. Samples taken from peak Activin A expression times are shown (96 hpi for conditions 1-5, 72 hpi for condition 6). E. Multi-step purification of mature Activin A from conditioned media from insect cells infected with DefBac^Fur+^::Activin A. Samples were resolved on an SDSPAGE gel and stained with coomassie blue. Samples 1-5 were reduced with DTT prior to loading. F. To assess activity, purified mature Activin A was added to epiblast derived stem cells (EpiSCs) cultured on fibronectin. Fold changes in gene expression, normalized to beta-actin, were determined over 11 days by quantitative polymerase chain reaction (qPCR) for the pluripotency makers Oct4, Nanog, Fgf5, and for the lineage marker, Pax6 (mean +/- S.D.; n = 3 separate experiments). Activin A from a commercial source was used as a control (grey box).

To demonstrate that GFP expression was associated with the production of an infectious virus, we collected the conditioned media from transfected cells and diluted it into fresh cultures. After five days, these cultures were incubated with fluorescently-tagged antibody that recognizes the baculovirus GP64 coat protein on the plasma membrane of infected insect cells and then single cell FACS sorted. These singly sorted cells were then added to fresh cultures in a 96-well plate. Out of 96 wells, 34 wells contained clearly infected cells that also strongly expressed GFP. The remaining 62 wells showed neither signs of infection nor GFP expression. These results show that rescue was both precise and efficient and that only recombinant virus expressing the transgene supplied by the shuttle vector are replication-competent. This design ensures that the parental bacmid lacking the gene of interest will not produce functional baculovirus.

#### Deletion of two non-essential baculovirus genes improves protein expression

Proteins required for baculoviral host liquefaction are not necessary for infection in cell culture and can often interfere with target protein expression. Two examples include cathepsin (*v-cath*) and chitinase (*chiA*), which are abundantly secreted late in the lytic cycle. Kaba et al. (2004) demonstrated that deletion of these genes improved the integrity and yield of a secreted target protein, p67. To improve our system, we eliminated these genes in H092. The resulting bacmid, called DefBac, produced infective baculovirus when co-tranfected with a pOCC shuttle vector and enhanced protein expression of the TGF-β member Activin A (Figure 2A; see Applications section).

#### A versatile shuttle vector set enables combinatorial appending of tags to the target protein

It is often desirable to append different tags to the target protein to improve expression levels, solubility, purity and visualization. To facilitate this optimization process, we developed a compatible set of 146 shuttle plasmids, called Optimized Classic Cloning (pOCC) vectors, designed to recombine with DefBac and express the user’s GOI (Figure 1B). Using this system, one GOI can be easily inserted into numerous vectors to produce target proteins with various combinations of cleavable N- and C-terminal tags (Figure 1C). The Defbac viral genome and the most popular 54 pOCC shuttle vectors are available at www.addgene.org (Table 2). The remaining pOCC shuttle vectors are available upon request (see Table 1 for the complete collection).

Each pOCC vector contains the same cloning cassette demarcated by NotI and AscI restriction enzyme sites. This cassette encodes a constitutive promoter (gb3) driving the toxic *ccdB* gene (Figure 1B). A pOCC vector with no insert will kill ccdB-sensitive bacteria (e.g. DH5alpha) and therefore must be maintained in a resistant strain such as DB3.1. When the target GOI is successfully inserted between the NotI and AscI sites, the *ccdB* gene is lost and the plasmid can be maintained in ccdB-sensitive cells. This feature dramatically reduces isolation of vectors lacking insert (data not shown). If target GOIs contain internal sites for NotI and AscI enzymes, their removal by silent mutation is required. However, recognition sites for both enzymes are 8 bp in length; thus, in practice, this additional mutagenesis step is rarely needed.

On the 5’ and 3’ ends of the cloning cassette are modules encoding purification tags, fluorescent probes, conjugating tags, trafficking peptides, and cleavable linkers (Figure 1C, Table 1). The translational reading frame is adjusted so that the NotI and AscI cloning sites encode amino acids with small side chains often found in natural peptide linkers. Upstream of these sequences lie a polyhedrin promoter and a translational start site, while stop codons and transcriptional terminator elements lie downstream (Figure 1B). Once a GOI is synthesized with compatible NotI and AscI ends, it can be inserted or subcloned into pOCC vectors to produce in-frame fusion proteins.

#### An enhanced version of DefBac that highly expresses a pro-protein convertase

Many secreted proteins require pro-protein convertases to generate the functional, mature form. We hypothesized that high expression of such target proteins might overwhelm the endogenous proteolytic processing enzymes in the Golgi apparatus. To elevate furin protein levels, we replaced the conotoxin gene in DefBac with full-length cDNA encoding murine furin driven by the p6.9 promoter. This promoter directs transcription earlier during the lytic cycle compared to the strong polyhedrin (polh) promoter driving target gene expression (Hill-Perkins and Possee, 1990). Thus, increased levels of furin are present before the target is translated. We call this parental bacmid “DefBac^Fur+^”.

### Applications

#### Expression and secretion of mature TGF-β family proteins

Transforming growth factor (TGF) Beta family proteins are secreted proteins involved in inter-cellular communication during development and tissue homeostasis. After translation, these proteins are imported into the ER in a pro-form, then trafficked to the Golgi apparatus, where they undergo maturation through proteolytic cleavage by furin convertases. The mature form is then secreted to the extracellular space by exocytosis. Recombinant TGF-β proteins are difficult to express at high levels in baculovirus-infected insect cells due to limited capacity of the secretory pathway. To overcome this limitation and improve expression of mature TGF-β family proteins, we employed our enhanced version of DefBac, called DefBac^Fur+^, which expresses furin.

We focused on production of recombinant Activin A, a secreted TGF-β protein involved in inflammation, neural development, and hematopoiesis (Chen et al., 2006; Vale et al., 1986). We co-transfected insect cells with a pOCC plasmid (pOCC-RL) containing Activin A and parental bacmid, then collected conditioned media containing secreted protein for 6 days post infection (hpi). We achieved higher Activin A yields using the DefBac parental bacmid compared to a conventional bacmid used in commercially-available systems (bMONH092, the bacmid encoding the baculvirus genome in the Bac-to-Bac system) (Figure 2A). This improvement is due to the elimination of genes encoding cathepsin and chitinase, two secreted proteins that are only required for host liquefaction (Figure 2B). However, the majority of Activin A was in the inactive pro-form (55 kDa); only trace amounts of the active, mature Activin A form (13 kDa) appeared (Figure 2A,B; reducing conditions).

We next sought to improve production yields of mature Activin A. To become fully active, TGF-β-family proteins require C-terminal cleavage by furin convertases that reside in the early Golgi apparatus and recognize the multi-basic motif RX(K/R)R (Thomas, 2002). Western blot analysis revealed that DefBac^Fur+^ greatly improved the production of mature Activin A and its secretion into the extracellular medium, with levels peaking 120 hpi (Figure 2C). We detected only trace amounts of the pro-form of Activin A in the DefBac^Fur+^ sample (Figure 2C). DefBac^Fur+^ also produced more mature Activin A compared to cultures co-infected with virus expressing Activin A and virus expressing furin (Figure 2D).

Next, we purified this mature Activin A and determined its activity. We isolated Activin A from the conditioned medium by cation exchange chromatography (SP), followed by hydrophobic interaction (HIC) and size exclusion chromatography (Superdex 75). From 1 L of conditioned medium, 0.5 mg of pure Activin A was obtained (Figure 2E). Addition of purified Activin A (from 10-40 ng/mL) to epiblast-derived stem cells (epiSC) suppressed expression of the differentiation marker Pax6 and either maintained or potently promoted the expression of pluripotency markers such FGF5, Nanog, and Oct4, indicating that Activin A was active (Figure 2F).

The TGF-β family represents a broad class of proteins that include growth, differentiation, and morphogenetic factors. In addition to Activin A, two prominent examples are human growth/differentiation factor 15 (GDF15) and human bone morphogenetic protein 4 (BMP4) (Wakefield and Hill, 2013). Expression of these targets using DefBac^Fur+^ greatly improved their processing and secretion levels (Figure 3A and 3B). Thus, DefBac^Fur+^ is generally suitable for recombinant production of active forms of secreted TGF-β proteins.

**Figure 3.**
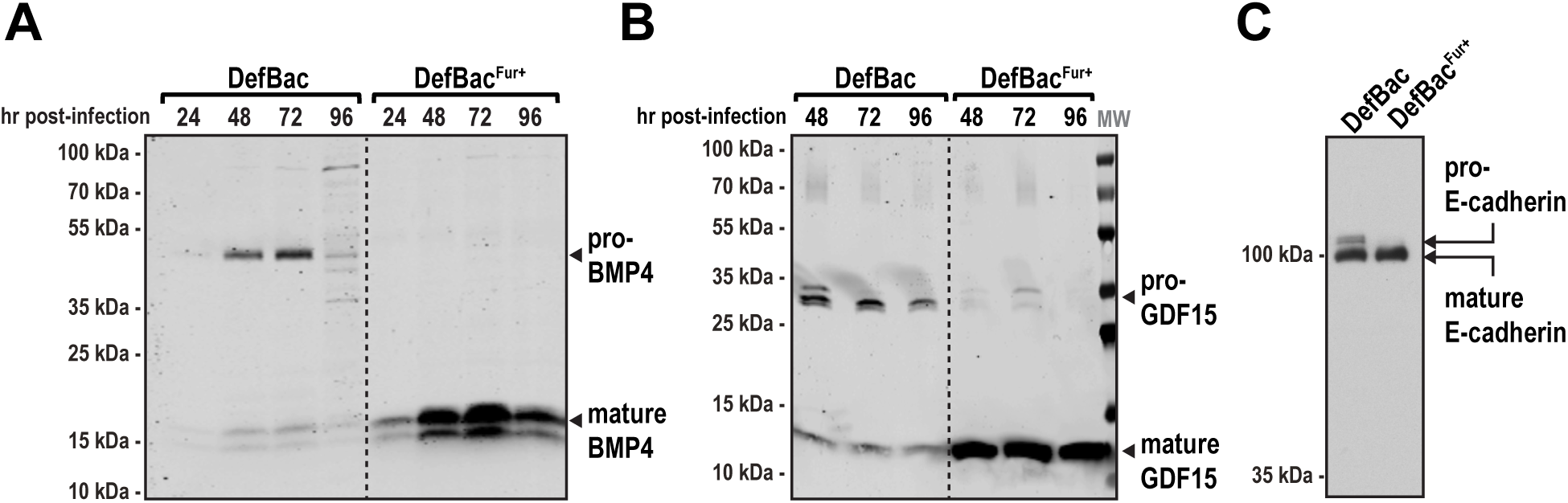
Production of mature BMP4, GDF15, and E-cadherin using DefBac^Fur+^. A. Human Bone morphogenetic protein 4 (BMP4) was expressed in insect cells using DefBac and DefBac^Fur+^ expression systems. BMP4 levels in the conditioned media were analyzed by western blot using an anti-BMP4 antibody. B. Human Growth/differentiation factor 15 (GDF15) was expressed in insect cells using DefBac and DefBac^Fur+^. GDF15 levels in the conditioned media were analyzed by western blot using an anti-GDF-15 antibody. C. The ecto-domain of canine E-cadherin was expressed in insect cells using DefBac and DefBac^Fur+^. E-cadherin levels in the conditioned media were analyzed by western blot using an anti-E-cadherin antibody.

#### Expression and secretion of processed E-cadherin

To test whether other secretory protein families similarly benefit from co-expression with furin, we generated recombinant baculovirus to express the ectodomain of canine E-cadherin, which requires maturation of the pro-form by a furin-like convertase. When the E-cadherin ectodomain is produced from DefBac virus, some unprocessed, pro-form is secreted into the conditioned medium (Figure 3C, lane 1) However, when we used DefBac^Fur+^, only the mature, fully-processed ectodomain was secreted (Figure 3C, lane 2). We conclude that DefBac^Fur+^ is suitable for production and secretion of a range of secreted proteins that require furin-dependent processing.

#### Expression of elongated, coiled-coil centrosome proteins

Centrosomes are micron-scale organelles that nucleate and organize microtubules needed for mitotic spindle assembly. Centrosomes comprise barrel-shaped centrioles surrounded by a less structured mass of protein called pericentriolar material (PCM). Mechanistic understanding of how PCM assembles and nucleates microtubules has long been hampered by lack of *in vitro* reconstitution approaches. This is attributed to the PCM scaffold being composed of elongated, high-molecular weight proteins that contain coiled-coil domains, which have been notoriously difficult to reconstitute in their full length.

The main PCM scaffold protein in the nematode *C. elegans*, SPD-5, contains 9 predicted coiled-coil domains, has a molecular weight of 135 kDa, and a hydrodynamic radius of ~8 nm (Hamill et al., 2002; Woodruff et al., 2017). Compared to the Bac-to-Bac (ThermoFisher) system, use of the FlexiBAC system improved the consistency and yield of full-length SPD-5 (Figure 4A). We then used FlexiBAC to express and purify 12 additional full-length PCM proteins (Figure 4B), which were sufficient to assemble minimal PCM and nucleate microtubule asters *in vitro* (Woodruff et al., 2017). Thus, the FlexiBAC system is suited for expressing elongated coiled-coil proteins, as well as globular kinases and microtubule-stabilizing enzymes.

**Figure 4.**
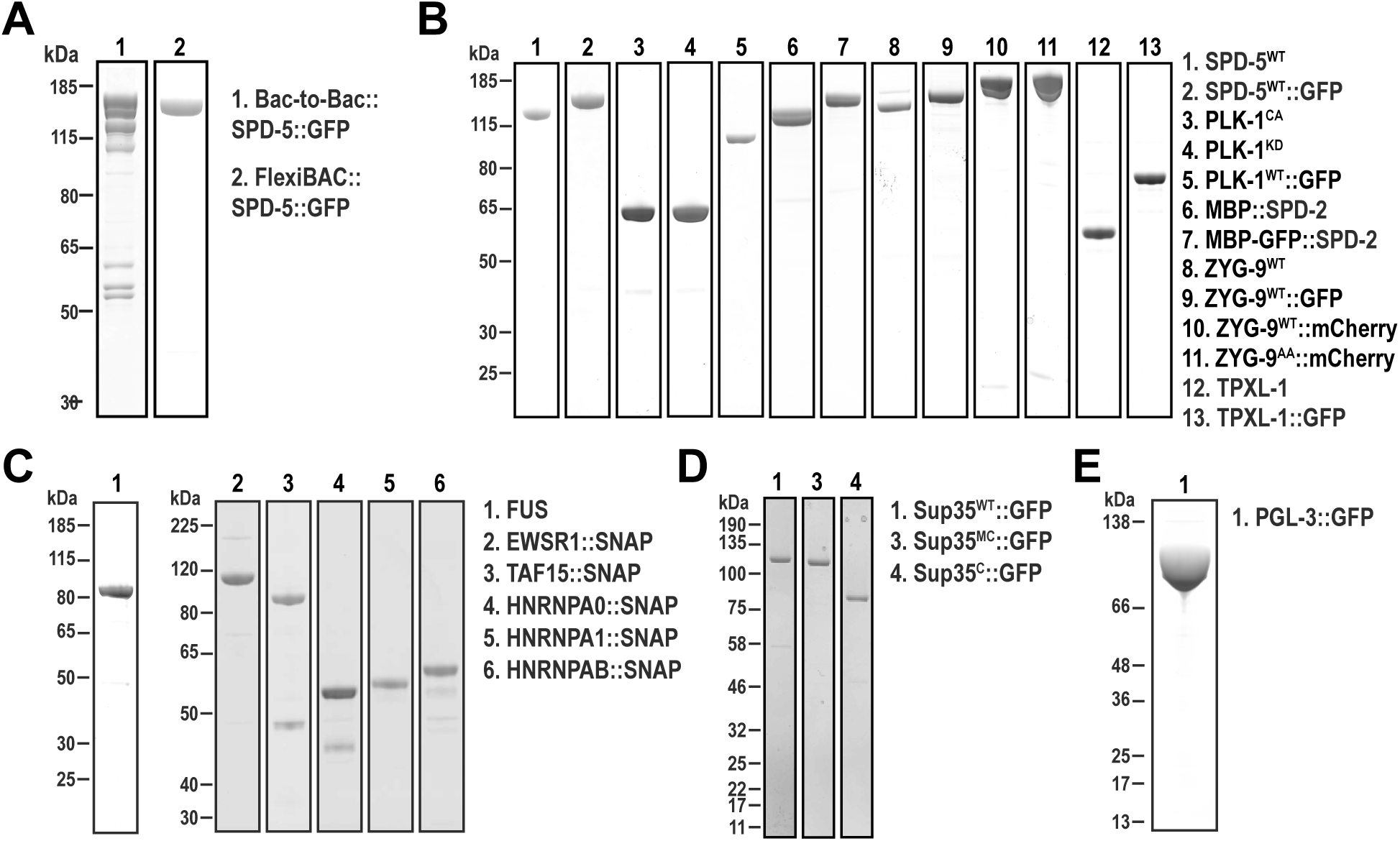
Production of recombinant intrinsically-disordered and centrosome proteins using DefBac. A. Coomassie-stained gels depicting 6xHis-tagged SPD-5::GFP expressed with the Bac-to-Bac system (ThermoFisher) or with FlexiBAC. In both lanes, SPD-5::GFP was purified from insect cell lysate using Ni-NTA affinity chromatography, then eluted with 250 mM imidazole. B. All proteins in (B-E) were expressed in SF9 insect cells using the FlexiBAC system. Coomassie-stained gels depicting purified *C. elegans* centrosome proteins. For more information, see (Woodruff et al., 2017; Woodruff et al., 2015). C. Coomassie-stained gels of purified human proteins that contain long stretches of intrinsically-disordered regions. For more information, see (Patel et al., 2015; Wang et al., 2018). D. Coomassie-stained gels depicting purified yeast prion-like proteins. MC and C represent middle and c-terminal domains of Sup35. For more information, see (Franzmann et al., 2018). E. Coomassie-stained gel of purified PGL-3, a protein that constitutes P granules in *C. elegans* embryos. For more information, see (Saha et al., 2016).

#### Expression of intrinsically-disordered proteins that self-assemble

Approximately 20% of the proteome in eukaryotes comprises protein sequences that lack globular structure, termed “intrinsically-disordered” proteins (IDPs)(Peng et al., 2015). Numerous non-membrane-bound organelles—such as nucleoli, stress granules, and germ granules—assemble through the coalescence of IDPs through multivalent interactions with themselves, other IDPs, or RNAs (Banani et al., 2017; Hyman and Brangwynne, 2011). Historically, IDPs have been difficult to reconstitute, as they are prone to aggregation and end up in inclusion bodies.

The FlexiBAC system was successful in expressing full-length IDPs that form the basis of stress granules (in human cells and yeast) and germ granules (in *C. elegans*) (Figure 4C-E). Target proteins were isolated from infected cells after multi-step purifications in sufficient yields (1-100 mg/L culture) to carry out extensive biochemical reconstitution assays. Under specific buffer conditions, each of these proteins self-assembles into micron-scale condensates (Franzmann et al., 2018; Patel et al., 2015; Saha et al., 2016) (Wang et al., 2018). We conclude that FlexiBAC is suitable for expression of disordered proteins that exhibit a strong tendency to self-assemble.

#### Discussion

Baculovirus expression has long been used for generating high yields of eukaryotic proteins that are properly folded and have native post-translational modifications. To make baculovirus expression simpler, more reliable, and more flexible, we generated a new system called FlexiBAC. We have demonstrated that FlexiBAC can be used to express diverse targets, including secreted proteins that require proteolytic processing in the Golgi apparatus, as well as intrinsically disordered and coiled-coil proteins.

FlexiBAC exhibits a number of benefits. First, the time from initial cloning to protein production can be as little as 13 days. Second, FlexiBAC is versatile and amenable for screening, as we designed a compatible shuttle vector set (146 vectors) harboring numerous tags used for purification, visualization, trafficking, and labeling with chemical probes. The target gene of interest is easily swapped between the plasmids using classic restriction enzyme cloning techniques. Third, FlexiBAC is suitable for expression of secreted proteins and is ideal for proteins requiring processing by the Golgi-resident furin convertase. Using a viral backbone that overexpresses furin, called DefBac^Fur+^, we produced high yields of Activin A, a TGF-β family protein. We expect our system to be useful for expressing mature versions of other targets requiring processing by a pro-protein convertase. Finally, the FlexiBAC system is open source and freely available through www.addgene.org.

FlexiBAC has been useful for expressing difficult proteins in numerous studies (Boke et al., 2016; Franzmann et al., 2018; Hernandez-Vega et al., 2017; Murray et al., 2016; Patel et al., 2015; Saha et al., 2016; Wang et al., 2018; Woodruff et al., 2017; Woodruff and Hyman, 2015; Woodruff et al., 2015). We envision additional modifications to specifically address other classes of protein targets. For example, with proteins found in heteromeric complexes, expression of individual subunits often is unsuccessful but is greatly improved by stoichiometric co-expression of all the subunits. We are currently adapting the FlexiBAC system to co-express multiple genes for stoichiometric expression of all subunits (Weissmann et al., 2016). In addition, FlexiBAC could also be expanded to express additional factors necessary for native post-translational modifications such as other pro-protein convertases, kinases, phosphatases, and glycosylation enzymes.

The speed, simplicity, and flexibility of the FlexiBAC system make it an ideal starting platform for expression and screening of diverse types of recombinant proteins. Given that FlexiBAC is open source, we invite improvements that will expand its applicability and facilitate production of a broader range of proteins.

## MATERIALS AND METHODS

Additional information about construction of the FlexiBAC plasmids and viral backbone can be found in the supplement.

### FlexiBAC Protocol

#### Reagents

- 24-well round-bottom plates (Qiagen #19583)
- SF9-ESF insect cells (Expression Systems # 94-001F)
- ESF921 Insect cell culture medium (Expression Systems #96-001-01)
- Autoclaved 2mL tubes
- Breathe-Easy sealing membranes (Sigma #Z380059-1PAK)
- 0.45 um PVDF syringe filters
- 10 ml and 60 ml syringes
- Pen-Strep (Gibco)
- Fetal Bovine Serum (Gibco)
- Escort IV transfection reagent (Sigma #L-3287)
- DNA isolation kit (Macherey-Nagel; Nucleobond Xtra Maxi 740414.50)
- SbfI-HF (NEB R3642-S)

#### Prep Phase 1. Make a stock of the DefBac viral backbone

1. Inoculate 1 L LB media with DefBac transformants (in DH10B cells). Add 30 mg Kanamycin (30 μg/ml final concentration). Incubate overnight until stationary phase.
2. Harvest bacteria, split into two tubes. Isolate DefBac using a maxi prep DNA isolation kit. Stop at the 70% ethanol wash step.
3. IMPORTANT: from here on, work in a sterile hood. Decant the final 70% ethanol wash. Do not dislodge the DNA pellet.
4. Air dry DNA (5-10 min). Do not over-dry the pellet.
5. Reconstitute DNA with 500 μl TE (sterile). Just add the buffer and let the DNA dissolve overnight at 4°C. Then mix by gentle pipetting (you want to avoid shearing).
6. Measure concentration of DNA by determining the absorbance at 260 nm (e.g. using a NanoDrop).
7. Digest with SbfI-HF enzyme to linearize DefBac. Incubate 3 hr at 37°C, then 20 min at 65°C to inactivate the SbfI.

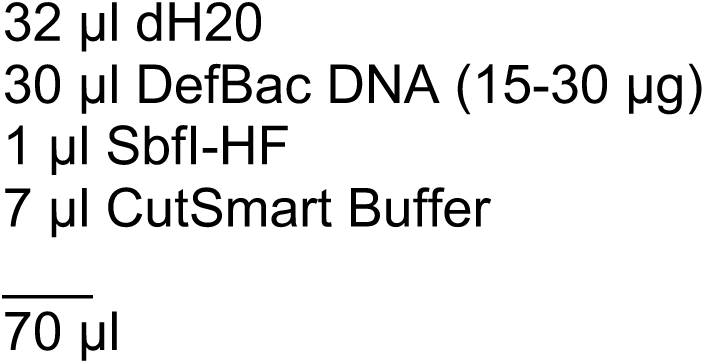
8. Determine the proper amount of the linearized DefBac DNA using transfection with a control plasmid (pOCC5; this expresses eGFP). Add 4, 2,1 0.5, 0.25 μL of SbfI-digested DefBac DNA. Follow steps described in Productions Phase 1, Day 1.

#### Prep Phase 2. Subclone target gene of interest into pOCC vector and miniprep

1. Use NotI and AscI restriction enzymes to digest pOCC vector and DNA encoding the target gene of interest.
2. Ligate plasmid and inserts, transform into DH5alpha cells, and grow cells overnight on LB + ampicillin plates.
3. Pick positive transformants and grow overnight cultures. Isolate DNA using a mini prep kit.

#### Production Phase 1. Transfect SF9 cells with pOCC plasmid and DefBac viral backbone to make the P1 virus stock

*A few days before transfection:*

Grow up an appropriate amount of SF9 insect cells. We typically passage cells on Friday morning to 0.5 × 10^6^ cells/ml and let them grow to 4-5 × 10^6^ cells/ml, which is usually by Monday morning. This culture is directly used for transfection.

*Day 1:*

1. Spray the 24-well plates with 70% ethanol and air dry in the hood.
2. Prepare transfection mix. In each well add reagents in the following order:

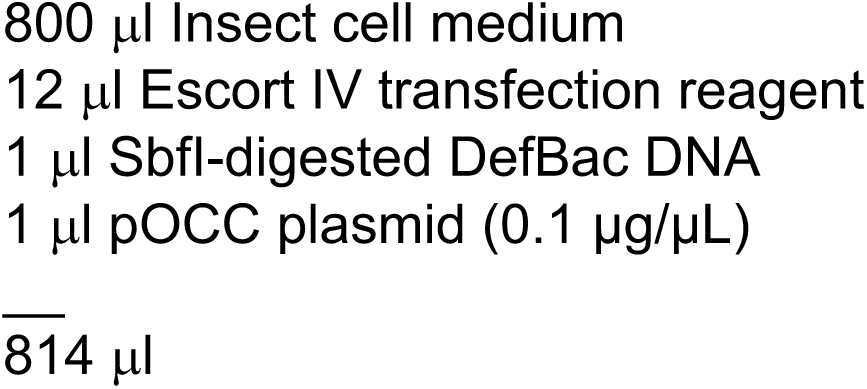
3. Gently mix the plate by shaking and incubate under the hood for 15 min. Add 200 μl cells per well (5 × 10^6^ cells/ml stock; this will result in 1 × 10^6^ cells/ml cells in the final mix).
4. Cover plate with a Breathe-Easy membrane.
5. Seal membrane/plate boundary using parafilm.
6. Place the sealed plate in an incubated shaker (200 rpm, 27°C).

*Day 2:*

1. Make master mix of insect cell medium + 1:50 Pen-Strep + 4% Fetal Bovine Serum. (Note: addition of serum is optional but is advised for long term storage)

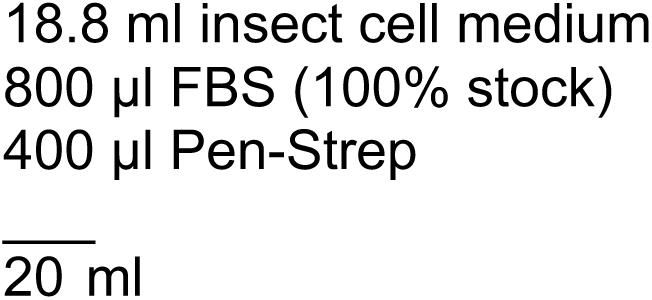
2. Take out the sealed plate from the incubated shaker. In a sterile hood, add 1 ml of master mix to each well that contains insect cells. This will result in final concentrations of 1% Pen-Strep and 2% FBS.
3. Cover plate with Breathe-Easy membrane and seal with the edges of the membrane with parafilm. Place the sealed plate in an incubated shaker (200 rpm, 27°C).

*Day 5:*

1. Gently spin the plates at 500xg for 5 min at room temperature.
2. Recover supernatant (1.5 ml). This contains released virus, called P1 virus.
3. Store P1 virus at 4°C protected from light. The P1 virus can also be frozen using a cryoprotectant in a slow cooling device as per freezing cultured cells.

**Production Phase 2. Amplify virus to get P2 stock**.

1. A few days prior, prepare an appropriate amount of insect cells so that they will be in log phase and at a density of 0.5 × 10^6^ cells/ml on the day of infection.
2. Add 50 μl P1 virus to 50 ml cells in a 250 ml flask. Add FBS (2% final concentration) and Pen-Strep (1% final concentration). (Note: addition of serum is optional but is advised for long term storage)
3. Shake for 5 days in an incubated shaker (100 rpm, 27°C).
4. Spin down cells at 2000xg for 5 min at room temperature. Collect supernatant and pass through a syringe filter. This is the P2 virus stock.
5. Store P2 virus at 4°C protected from light. The P2 should be good for ~1 month.

**Production Phase 3. Express target protein and harvest cells**.

1. Infect log phase insect cells (1 × 10^6^ cells/ml) with virus. In the beginning, try a 1:100 dilution of P2 virus; e.g., add 5 ml virus for 500 ml insect cells. If optimization is necessary, try a dilution series.
2. Shake infected cells for 72 hr (100 rpm, 27°C). Note: to determine the optimal time of harvesting, we advise performing a time course of expression. Collect samples at 48, 72, 96 hours post infection and determine the level of expression of the target protein by SDS-PAGE and western blot.
3. Harvest cells using a gentle spin and a slow deceleration (300xg for 10 min).
4. Resuspend cell pellet with an appropriate buffer (no detergent!) and transfer to several 14 ml Falcon tubes.
5. Flash freeze Falcon tubes in liquid nitrogen. Store at −80°C. Note: some proteins are prone to degradation after freeze/thaw. In this case, it is advisable to proceed with extract preparation and purification using the fresh cell pellet.

## ACKNOWLEDGEMENTS

J.B.W. is supported by the Cancer Prevention Research Institute of Texas (CPRIT grant RR170063) and the Endowed Scholars program at UT Southwestern. D.N.D was supported by the Max Planck Society.

